# Spatiotemporal dynamics of PIEZO1 localization controls keratinocyte migration during wound healing

**DOI:** 10.1101/2020.10.18.344598

**Authors:** Jesse R. Holt, Wei-Zheng Zeng, Elizabeth L. Evans, Seung-Hyun Woo, Shang Ma, Hamid Abuwarda, Meaghan Loud, Ardem Patapoutian, Medha M. Pathak

## Abstract

Keratinocytes, the predominant cell type of the epidermis, migrate to reinstate the epithelial barrier during wound healing. Mechanical cues are known to regulate keratinocyte re-epithelization and wound healing however, the underlying molecular transducers and biophysical mechanisms remain elusive. Here, we show through molecular, cellular and organismal studies that the mechanically-activated ion channel PIEZO1 regulates keratinocyte migration and wound healing. Epidermal-specific *Piezo1* knockout mice exhibited faster wound closure while gain-of-function mice displayed slower wound closure compared to littermate controls. By imaging the spatiotemporal localization dynamics of endogenous PIEZO1 channels we find that channel enrichment in sub-cellular regions induces a localized cellular retraction that slows keratinocyte migration. Our findings suggest a potential pharmacological target for wound treatment. More broadly, we show that nanoscale spatiotemporal dynamics of Piezo1 channels can control tissue-scale events, a finding with implications beyond wound healing to processes as diverse as development, homeostasis, disease and repair.

## Introduction

The skin, the largest organ of the body, serves as a barrier against a myriad of external insults while also performing important sensory and homeostatic functions. Cutaneous wounds interfere with all these functions and expose the body to an increased risk of infection, disease and scar formation (Evans et al. 2013). During the repair of wounded skin, the migration of keratinocytes from the wound edge into the wound bed plays an essential step in re-establishing the epithelial barrier and restoring its protective functions (Kirfel and Herzog 2004; Gantwerker and Hom 2011). Accumulating evidence has shown that mechanical cues and cell-generated traction forces in keratinocytes play an important role in regulating the healing process and wound closure (Evans et al. 2013; Rosińczuk et al. 2016; Brugués et al. 2014-9; Hiroyasu, Colburn, and Jones 2016-6; Huang, Du, and Ogawa 2017; Ladoux and Mège 2017). However, the molecular identity of keratinocyte mechanotransducers that control re-epithelialization remains unknown. Cells are able to sense and detect mechanical forces, converting them into biochemical signals through the process of mechanotransduction. One class of mechanosensors utilized by cells are mechanically-activated ion channels which offer the unique ability for cells to rapidly detect and transduce mechanical forces into electrochemical signals (Nourse and Pathak 2017; Murthy, Dubin, and Patapoutian 2017). The Piezo1 ion channel has been shown to play an important role in a variety of cell types, and it regulates several key biological processes including vascular and lymphatic development, red blood cell volume regulation, stem cell fate, the baroreceptor response, cardiovascular homeostasis, cartilage mechanics, and others (J. Li et al. 2014; Ranade et al. 2014; Pathak et al. 2014; Rocio Servin-Vences et al. 2017; Zeng et al. 2018; Nonomura et al. 2018; Cahalan et al. 2015; W. Lee et al. 2014). Previous studies in MDCK cells and in zebrafish larvae have demonstrated the importance of the channel in homeostatic regulation of epithelial cell numbers (Gudipaty et al. 03 02, 2017; Eisenhoffer et al. 2012). As yet, the role of Piezo1 in skin wound healing, an important epithelial function, has not been investigated. We asked whether Piezo1 may function as a mechanosensor regulating keratinocyte re-epithelialization during the wound healing process. Here we show that Piezo1 activity reduces efficiency of keratinocyte migration and wound healing, and inhibition of Piezo1 results in faster wound healing *in vitro* and *in vivo*. It achieves this via dynamic changes in its subcellular localization, concentrating at areas of the wound edge and causing local retraction at these regions.

## Results

### Reduced PIEZO1 accelerates wound healing

Analysis of *Piezo* channel mRNA expression in mouse tissues has previously shown that *Piezo1* is highly expressed in skin, while *Piezo2* is less abundant (Coste et al. 2010). To characterize PIEZO1 expression profile in skin, we used a reporter mouse expressing a promoter-less β-geo (β-gal and neomycin phosphotransferase) in-frame with a portion of the PIEZO1 channel (Ranade et al. 2014). LacZ staining of skin tissue from these reporter mice revealed a high expression of PIEZO1 in the epidermal layer of keratinocytes as well as in hair follicles (Fig. 1A). Since the global knockout of *Piezo1* is embryonically lethal (Ranade et al. 2014; J. Li et al. 2014), we generated an epidermal-specific knockout mouse to investigate whether PIEZO1 plays a role in cutaneous wound healing. The *Krt14Cre* mouse line was crossed with *Piezo1*^*fl/fl*^ mice (Cahalan et al. 2015) to generate *Krt14Cre;Piezo1*^*fl/fl*^ mice (hereafter referred to as conditional knockout, cKO) which are viable and develop normally (Supplementary Fig. 1), consistent with observations by Moehring et al (Moehring et al. 2020). qRT-PCR analysis using keratinocytes harvested from *Piezo1* cKO and littermate control animals confirmed the expression of *Piezo1* but not *Piezo2* in control mice (Supplementary Fig. 2), and showed that *Piezo1* mRNA expression is efficiently abrogated in cells from cKO animals (Fig. 1B). Furthermore, we also generated the *Krt14Cre;Piezo1*^*cx/+*^ mouse line which expresses the gain of function (GoF) *Piezo1* mutation, R2482H (Ma et al. 2018), in keratinocytes.

**Figure 1.**
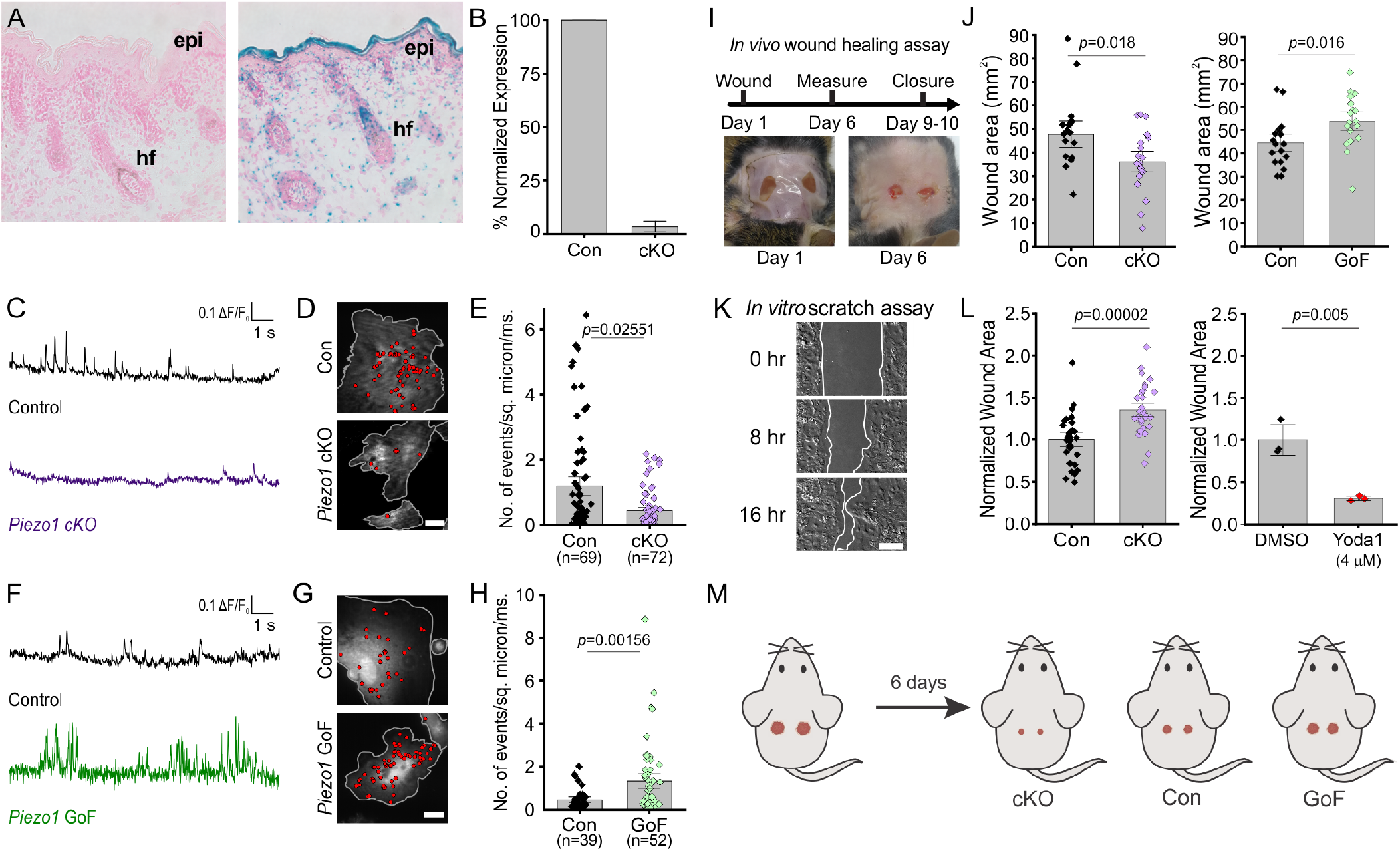
PIEZO1 is expressed in keratinocytes, produces Ca^2+^ flickers and regulates skin wound healing. **(A)** Representative images of LacZ stained Piezo1^+/+^ (*left*) and Piezo^+/βGeo^ (*right*) skin sections from P2 (postnatal day 2) mice. epi, epidermis. hf, hair follicle. **(B)** qRT-PCR from primary neonatal keratinocytes of *Piezo1* mRNA expression in Krt14^+/+^ *Piezo1*^*fl/fl*^ (Con) and *Krt1*4^*Cre/+*^;*Piezo1*^*fl/fl*^ (cKO) mice. See also Supplementary Figures 1-2. Data collected from 2 litters. **(C)** Representative examples of Ca^2+^ flickers recorded from Control (*top)* and *Piezo1-*cKO (bottom) keratinocytes. Traces show fluorescence ratio changes (ΔF/F_0_) from a region of interest, plotted against time. **(D)** Representative images of sites of Ca^2+^ flickers (red dots) overlaid on images of keratinocytes isolated from control (*top*) and littermate *Piezo1-*cKO (*bottom*) mice that are loaded with fluorescent Ca^2+^ indicator Cal-520AM. Grey line denotes cell boundary, Scale Bar = 20 μm. **(E)** Frequency of Ca^2+^ flickers in *Piezo1-*cKO and respective control cells. n in E denotes the number of videos (i.e., unique fields of view imaged, each of which is composed of one or more cells). Videos were collected from 4 independent experiments from 3 litters. See also Supplementary Video 1. **(F, G, H)** Similar to C, D, E but using keratinocytes from control (*top*) and littermate *Krt14Cre;Piezo1^cx/+^* (GoF) (*bottom*) mice. Videos were collected from 4 independent experiments from 3 litters. See also Supplementary Video 2. **(I)** Diagram of *in vivo* wound healing model (*top*) and representative wound images at Day 0 and 6 (*bottom*). **(J)** Wound area of *Piezo1* cKO (*left*) and *Piezo1* GoF (*right*) groups at Day 6 relative to control (Con) littermates. Control (Con) refers to the Cre-negative littermates in each group (n = 18 wounds for Controls and *Piezo1* GoF, and n = 20 wounds for *Piezo1* cKO, with two wounds per animal). **(K)** Representative images of an *in vitro* scratch assay. White line represents the monolayer edge. Scale bar = 200 μm. **(L)** Quantification of scratch wound area (corresponding to area depicted between the two white lines in I, 16 hours after wound generation in Control vs littermate Piezo1-cKO keratinocytes (*left*); n = 29, WT, n = 31 Piezo1-cKO images from 3 independent experiments) or DMSO-treated vs 4 μM Yoda1 treated cells (*right*; n = 3 independent experiments). Data are normalized to the control condition. **(M)** Schematic illustrating results from *in vivo* wound healing assay shown in I and J. Mice were wounded (*left*) and after 6 days, wounds of *Piezo1-*cKO mice healed more than Control, whereas wounds from *Piezo1-*GoF mice healed less. All bars denote mean ± S.E.M.

To confirm functional change to PIEZO1 in mutant keratinocytes, we performed Ca^2+^ imaging using Total Internal Reflection Fluorescence Microscopy (TIRFM) of keratinocytes isolated from *Piezo1* cKO, GoF, and their respective control (Cre-) littermates. We previously reported that in adherent cells Piezo1 produces Ca^2+^ flickers in response to cell-generated forces in the absence of external mechanical stimulation (Pathak et al. 2014; Ellefsen et al. 2019). Compared to littermate control mice, keratinocytes from *Piezo1*-cKO mice showed a 63% reduction in Ca^2+^ flicker activity, indicating that a majority of Ca^2+^ flickers arise from PIEZO1 activity (Fig. 1C, 1D, 1E, Supplementary Video 1). On the other hand, *Piezo1*-GoF cells displayed a nearly threefold increase in flicker frequency relative to littermate controls demonstrating the mutation has the desired effect on channel activity in keratinocytes (Fig. 1F, 1G, 1H, Supplementary Video 2).

To investigate the function of PIEZO1 in keratinocytes *in vivo*, we generated full-thickness wounds on the dorsal skin of *Piezo1* cKO, GoF and their respective control littermates and assessed wound closure (Fig 1I). Six days post wounding, *Piezo1*-cKO mice displayed significantly smaller wound areas relative to their control littermates, while *Piezo1*-GoF mice showed larger wound areas, suggesting increased channel activity leads to impaired rates of wound closure (Fig 1J).

To determine that the effect on wound healing was caused by changes to the rate of keratinocyte reepithelialization, we mimicked the *in vivo* wound healing paradigm, *in vitro*. We generated scratch wounds in keratinocyte monolayers to trigger the re-epithelialization process, and allowed the monolayers to migrate toward each other (Fig. 1K). Scratches in monolayers of *Piezo1*-cKO keratinocytes closed faster than those from littermate control cells (Fig. 1L, left). Conversely, when the Piezo1 agonist Yoda1 was added to healing control monolayers, scratch wound closure was significantly impaired (Fig. 1L, right), further suggesting PIEZO1 involvement in re-epithelialization. Collectively, our *in vitro* and *in vivo* data suggest that the PIEZO1 ion channel plays an important role in wound healing, with Piezo1 knockout accelerating wound healing (Fig. 1M).

### PIEZO1 activity regulates keratinocyte migration

To determine whether the differences in wound closure rates arise due to PIEZO1’s effect on keratinocyte motility during the re-epithelialization process, we captured migration dynamics of dissociated single keratinocytes from *Piezo1*-cKO mice. Isolated cells were sparsely seeded onto fibronectin-coated glass-bottom dishes and imaged over several hours using differential interference contrast (DIC) time-lapse imaging (Fig. 2A, Supplementary Video 3). We tracked the position of individual cells in the acquired movies and analyzed the extracted cell migration trajectories using an open-source algorithm, DiPer (Gorelik and Gautreau 2014). The time-lapse images, and corresponding cell migration trajectories (Fig. 2B; Supplementary Fig. 3) revealed that the migration patterns of *Piezo1*-cKO keratinocytes are distinct from their littermate control cells. To quantify cellular migration, we generated mean squared displacement (MSD) plots which provide a measure of the surface area explored by the cells, i.e. an indication of the overall efficiency of migration. Interestingly, *Piezo1*-cKO keratinocytes explored a larger area compared to littermate control cells (Fig. 2C).

**Figure 2.**
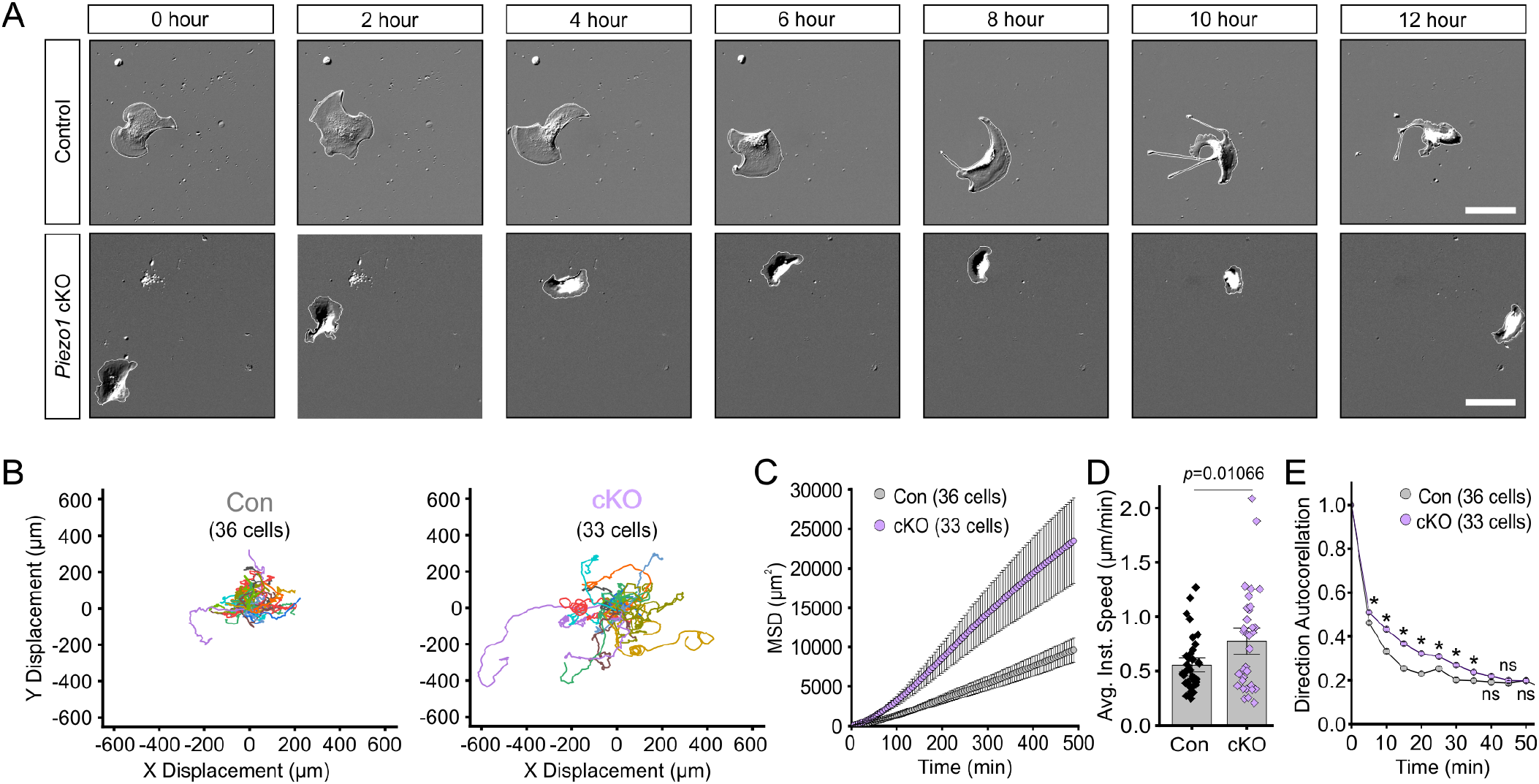
PIEZO1 mediates speed and direction of single cell keratinocyte migration. **(A)** Representative DIC images from time-lapse series of individual migrating keratinocytes isolated from Control (*top*) and respective *Piezo1-*cKO littermates (*bottom*). White lines denote the cell boundary. Scale Bar = 25 μm. **(B)** Cell trajectories derived from tracking keratinocytes during time lapse experiments. Trajectories are shown with cell position at time point 0 normalized to the origin. See also Supplementary Figure 3. **(C)** Mean squared displacement (MSD) analysis of control and *Piezo1-*cKO keratinocytes tracked in B. Average MSD is plotted as a function of time. Error bars (S.E.M) are smaller than symbols at some points. **(D)** Quantitation of the average instantaneous speed from individual *Piezo1-*cKO keratinocytes relative to littermate control cells. **(E)** Average direction autocorrelation measurement of *Piezo1-cKO* and control keratinocytes plotted as a function of time interval. * denotes a statistically significant difference, and ns denotes “not statistically significant”. From left to right: *p*= 2.0307×10^−4^, 5.75675×10^−14^, 3.18447×10^−15^, 5.34662×10^−10^, 1.72352 ×10^−4^, 1.34648 ×10^−5^, 0.01951, 0.13381, 0.61758 as determined by Kruskal-Wallis test. Plotted error bars (S.E.M) are smaller than symbols. Data are from 3 independent experiments from 2 litters. All bars denote mean ± S.E.M. See also Supplementary Video 3.

The MSD of a migrating cell is determined by two parameters: displacement rate (speed) and directional persistence (propensity of the cell to move in a straight line). The average instantaneous speed calculated by DiPer analysis was higher for *Piezo1*-cKO cells relative to littermate control cells (Fig. 2D). To assess directional persistence, we performed direction autocorrelation analysis, a robust measure of migration directionality that, unlike the more commonly used directionality ratio analysis, is not confounded by differences in migration speed (Gorelik and Gautreau 2014). The direction autocorrelation function for trajectories from *Piezo1*-cKO keratinocytes decayed slower than for littermate control cells, indicative of fewer turns (Fig. 2E). Thus, *Piezo1*-cKO keratinocytes migrate significantly faster and straighter. Overall, our data demonstrate that PIEZO1 regulates keratinocyte migration, with channel activity resulting in less efficient cellular migration.

### PIEZO1 activation causes cellular retraction

To determine how PIEZO1 may regulate keratinocyte migration behavior, we focused on cellular dynamics of single, migrating keratinocytes by imaging at a high spatiotemporal resolution. Using DIC imaging, we monitored migrating keratinocytes at 5 second intervals under control conditions and following Yoda1 treatment (Fig. 3A, Supplementary Video 4) and analyzed cell edge dynamics over time using kymographs to visualize position of the keratinocyte cell edge (Fig. 3B). We observed that under control conditions, the cell edge demonstrated cycles of protrusion and retraction which was expected since cell migration is known to progress by iterative cycles of protrusion and retraction (Giannone et al. 2004, 2007). PIEZO1 activation by 4 μM Yoda1 greatly affected these cycles and resulted in an extremely dynamic cell edge (Fig. 3B, Supplementary Video 4). The frequency as well as the velocity of cell edge retractions increased upon PIEZO1 activation and resulted in a net cellular retraction over time (Fig. 3B), with some cells demonstrating drastic retraction with Yoda1 treatment (Supplementary Video 5). *Piezo1*-cKO keratinocytes did not show an increase in retraction upon treatment with 4 μM Yoda1 (Fig. 3C, Supplementary Video 6), demonstrating that increased retraction is mediated by PIEZO1.

**Figure 3.**
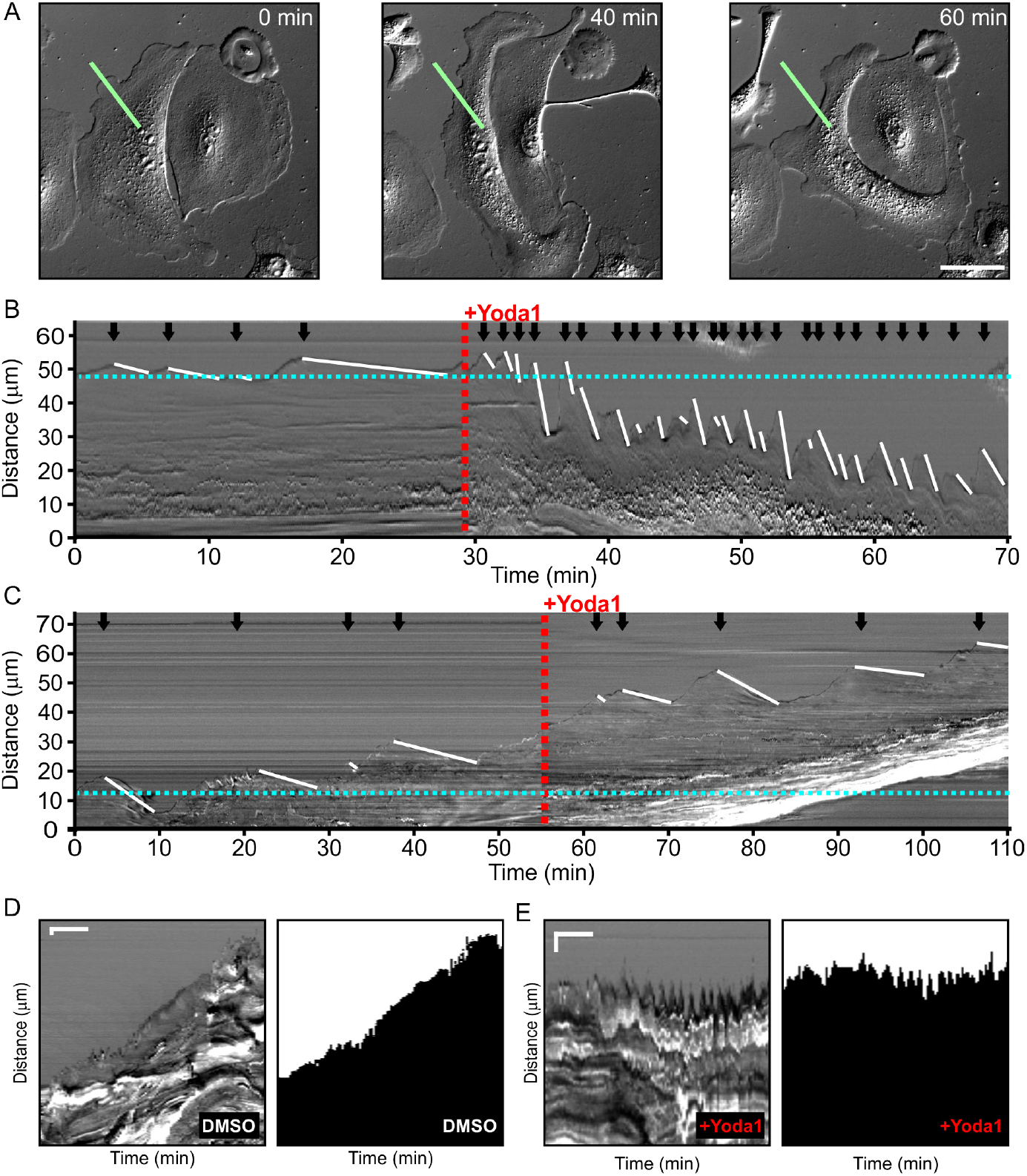
PIEZO1 activity promotes cellular retraction. **(A)** Images from a DIC time-lapse series of keratinocytes at time 0, 40 and 60 minutes. 4 μM Yoda1 was added to the bath solution after 30 minutes of baseline recording. The green lines denote the region of interest used for generating the kymograph in panel B. Scale Bar = 50 μm **(B)** Representative DIC kymograph of the keratinocyte presented in A. The red dotted line denotes the addition of 4 μM Yoda1. The black arrows indicate retraction events and the slope of the descending white lines denotes the velocity of retraction. Dotted blue line denotes starting position of the cell edge. See also Supplementary Video 4 and 5. **(C)** Representative DIC kymograph of a *Piezo1-*cKO keratinocyte with the same annotations as B. See also Supplementary Video 6. **(D)** Representative DIC kymograph taken at the wound edge, showing wound edge dynamics during an *in vitro* scratch assay in control (DMSO)-treated monolayers (*left*). Kymograph of the binarized version of the video, with the cell monolayer represented in black and cell-free space of the wounded area in white (*right*). Scale Bar = 10 μm. Time bar = 100 min. See also Supplementary Video 7. **(E)** As for D but in a sample treated with 4 μM Yoda1. The data in D and E are representative of 3 independent experiments from keratinocytes from 3 litters.

We then asked whether PIEZO1-mediated retraction events observed in single cells are relevant in the context of a wounded cell monolayer. We generated scratch wounds in keratinocyte monolayers and performed DIC time-lapse imaging of the healing monolayer in the presence and absence of 4 μM Yoda1 to observe dynamics of the wound edge. In control conditions, the monolayer advanced forward (Fig. 3D, Supplementary Video 7), while the presence of Yoda1 increased retraction events and prevented the monolayer from advancing far into the wound bed (Fig. 3E, Supplementary Video 7). Collectively, our results show that Piezo1 activity increases cellular retraction in keratinocytes, both in single cells and monolayers, which has a net effect on cell migration.

### Dynamic PIEZO1 localization informs retraction events to regulate wound healing

To understand how PIEZO1 mediates retraction events, we visualized localization of endogenous PIEZO1 in single keratinocytes harvested from a *Piezo1*-tdTomato fusion knock-in reporter mouse (Ranade et al. 2014). Using this model, we previously reported punctate membrane localization of endogenous PIEZO1-tdTomato channels in neural stem/progenitor cells and mouse embryonic fibroblasts (Ellefsen et al. 2019). Here, we similarly visualized PIEZO1 localization, first in fixed keratinocytes by immunostaining for tdTomato. In non-polarized keratinocytes without an obvious migratory morphology, we observed PIEZO1-tdTomato puncta distributed throughout the cell consistent with our previous experiments in other cell types (Fig. 4A, left). However, in keratinocytes that displayed an overt lamellipodium, a highly dynamic sheet-like organelle at the leading edge of migrating cells (Ridley 2011), PIEZO1-tdTomato localization displayed a gradient, with higher PIEZO1 levels at the trailing or rear end of the cell (Fig. 4A, right). To confirm this observation, we directly imaged endogenous PIEZO1-tdTomato’s subcellular localization in live, migrating keratinocytes using TIRFM imaging. By tracking the cells over time we confirmed that enrichment of PIEZO1-tdTomato puncta occurred at the rear end of the cell (Fig. 4B, Supplementary Video 8). Importantly, the rear of the cell is a site of marked cellular retraction by nonmuscle myosin II-mediated contractile forces (Petrie, Doyle, and Yamada 2009; Yam et al. 2007).

**Figure 4.**
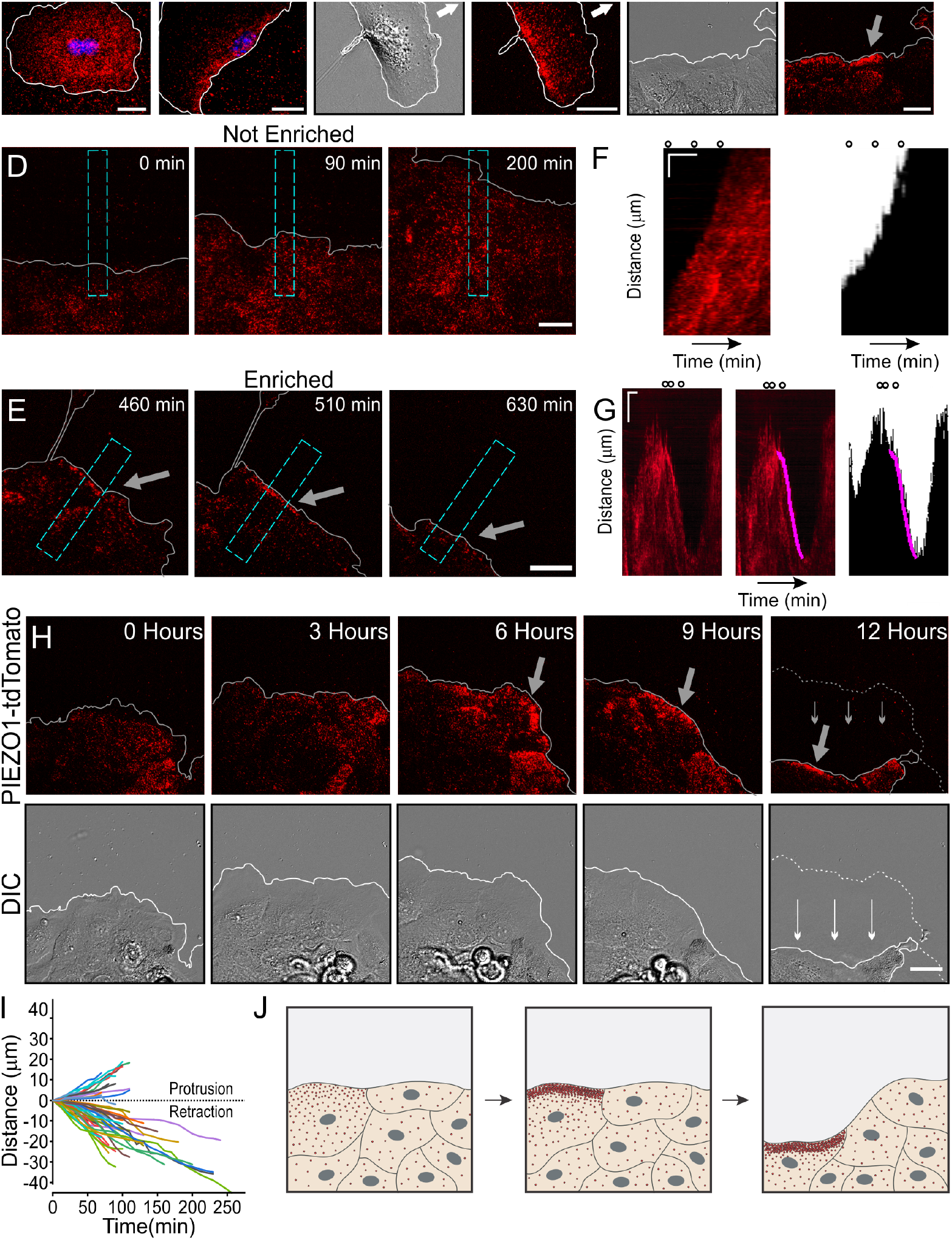
Dynamic PIEZO1 channel localization controls wound-edge retraction. Representative TIRFM images of a non-polarized PIEZO1-tdTomato keratinocyte (*left*) and polarized keratinocyte (right) immuno-labeled with an antibody against tdTomato (red) and nuclear stain Hoechst (blue). White lines denote the cell boundary. Scale Bar = 20 μm. **(B)** Representative DIC (*left*) and TIRF (*right)* images illustrating the location of PIEZO1-tdTomato protein in live migrating keratinocytes. White lines denote the cell boundary. Scale Bar = 20 μm. See also Supplementary Video 8. **(C)** Representative DIC (*left*) and TIRF (*right*) image visualizing the location of PIEZO1-tdTomato protein in live collectively migrating keratinocytes in an *in vitro* scratch assay. Grey lines denote the boundary of the cell monolayer. Grey arrow indicates region of PIEZO1 enrichment. Scale Bar = 20 μm. See also Supplementary Figure 5. **(D, E)** Representative TIRF images taken from time-lapse image series of healing monolayers from PIEZO1-tdTomato keratinocytes highlighting fields of view in which PIEZO1-tdTomato is not enriched **(D)** and enriched **(E)** at the monolayer’s leading edge. Grey lines denote the boundary of the cell monolayer. Arrow indicates regions of enrichment. Scale Bars = 20 μm. Blue dashed rectangles in D & E depict the regions used to generate kymographs in F and G. TIRF Images were acquired every 10 minutes over a period of 16.6 hours. **(F, G)** *Left panels*: Representative kymographs illustrating PIEZO1-tdTomato puncta dynamics during the time-lapse series shown in D and E, respectively. *Middle Panel (for G only):* Magenta line denotes periods of PIEZO1-tdTomato puncta enrichment at the wound edge, identified and tracked using the deep learning-based kymograph analysis software, Kymobutler. *(F)* No Kymobutler tracks were detected in non-enriched regions. *Right panels*: Representative kymographs from binarized versions of DIC images corresponding to D and E respectively, with the Kymobutler track output from the middle panel overlaid. The cell monolayer is represented in black, and white denotes cell-free space of the wounded area. Note the PIEZO1-tdTomato enrichment track correlates with period of cell retraction. Scale Bar = 10 μm. Time bar = 100 minutes. Black open circles on top represent the time-points of images shown in D and E. See also Supplementary Videos 9 and 10. **(H)** Representative DIC (*bottom*) and TIRF (*top*) images during a time-lapse imaging series following scratch generation at 0 hours showing PIEZO1-tdTomato localization and monolayer position. Grey (*top*) and white (*bottom*) lines denote the cell boundary. Dotted line in the 12-hour image denotes the position of the monolayer at 6 hours; small arrows indicate direction of monolayer movement during this period. Large grey arrow indicates region of PIEZO1 enrichment. Scale Bar = 20 μm. See also Supplementary Video 11. **(I)** Plot showing 54 individual PIEZO1-tdTomato Kymobutler tracks from 25 kymographs collected from 3 independent experiments after normalizing the starting spatial and time coordinates of each track to the origin. **(J)** Schematic of a healing monolayer indicating distributed Piezo1 localization (red dots) immediately post scratch (*left*), the development of areas of Piezo1 enrichment (*middle*) and subsequent retraction of those areas (*right*).

We then asked how PIEZO1 localization may mediate retraction events in wounded cell monolayers. We generated a scratch wound in a confluent monolayer of *Piezo1*-tdTomato keratinocytes and imaged spatiotemporal dynamics of PIEZO1-tdTomato localization at the cell-substrate interface using TIRFM imaging together with DIC imaging over a period of several hours. We found that in areas of the cell monolayer away from the wound edge, PIEZO1-tdTomato puncta showed no obvious enrichment, but at periodic regions along the wound margin we observed that PIEZO1 became enriched in band-like structures (Fig. 4C). Interestingly, this enrichment was highly dynamic, such that channel enrichment ebbed and flowed over the duration of imaging (Supplementary Videos 9 and 10).

To systematically assess the correlation of PIEZO1-tdTomato enrichment with wound edge retraction, we used kymographs to graphically represent PIEZO1-tdTomato position over the imaging period (Fig. 4F, 4G). PIEZO1-tdTomato enrichment events at the wound edge appeared as linear streaks in the kymographs (Fig. 4G, left panel) which could be objectively identified by Kymobutler, a deep-learning-based kymograph analysis software (Jakobs, Dimitracopoulos, and Franze 2019) (Fig. 4G, middle and right panels). For the kymographs from fields of view which did not show any apparent PIEZO1-tdTomato puncta enrichment near the wound edge (Fig. 4D, Supplementary Video 9), Kymobutler did not detect any tracks and we did not observe any retraction of the wound edge (Fig. 4F). In fields of view that exhibited channel enrichment at the wound edge (Fig. 4E, Supplementary Video 10), resulting tracks were overlaid on the DIC kymographs to examine migration dynamics of the wound edge in relation to Piezo1 enrichment (Fig. 4G). 72% of these tracks displayed a negative slope corresponding to cell edge retraction and aligned with retraction events (Fig. 4G, I). In some fields of view, PIEZO1 enrichment and the accompanying retraction lasted for shorter periods of time, and the periods without channel enrichment were accompanied by wound edge protrusion (Fig. 4D, F and Supplementary Video 9). In other cases, channel enrichment was maintained for several hours and was accompanied by a sustained and overt retraction of the wound edge throughout that period (Fig. 4H, Supplementary Video 11). Thus, enrichment of PIEZO1-tdTomato puncta resulted in wound edge retraction.

Taken together with the observations of PIEZO1-tdTomato dynamics in single cells, this analysis demonstrates that PIEZO1-tdTomato enrichment and activity induces cellular retraction. We propose that dynamic enrichment of the channel protein serves to amplify channel activity and the downstream retraction events. In wound-healing monolayers of keratinocytes, PIEZO1-tdTomato enrichment and the subsequent retraction observed at the wound edge (Fig. 4J) provides a molecular mechanism for how PIEZO1 delays wound healing, while absence of the channel accelerates wound healing.

## Discussion

Our findings demonstrate that epidermal-specific *Piezo1* knockout resulted in faster wound healing in mice, and conversely a *Piezo1* gain-of-function mutation slowed wound healing. We recapitulate this observation *in vitro,* and show through a combination of orthogonal assays in single cells and in monolayers that Piezo1 activity modulated by dynamic spatial reorganization of the channel protein slows keratinocyte cell migration during re-epithelialization. These findings provide physiological evidence for a mechanically-activated ion channel in wound healing and suggest potential therapeutics through a targeted inhibition of Piezo1, perhaps applied topically, that could help speed wound healing, potentially reducing risk of infection. Given that speedy wound healing affords an evolutionary advantage and that PIEZO1 activity slows wound healing, a puzzling question arises regarding the role of PIEZO1 expression in keratinocytes. Perhaps, there is an advantage to slower healing in the presence of PIEZO1, or the channel is important for other functions in keratinocytes. Consistent with this idea, a recent study by Moehring *et al.* reports that keratinocyte PIEZO1 is critical for sensory afferent firing and behavioral responses to innocuous and noxious mechanical stimulation (Moehring et al. 2020).

One of the most surprising findings to emerge from our studies is the highly dynamic nature of the spatial localization of PIEZO1 channels in migrating cells. Based on this finding, we propose a novel mechanism regulating cell migration wherein spatiotemporal enrichment of PIEZO1 channels serves to localize and amplify channel activity, and the resulting contractile force, to spatially control cellular retraction events. Piezo1 has been implicated in cell migration in different cell types *in vitro* (Maneshi et al. 2018; McHugh et al. 2012; Chubinskiy-Nadezhdin et al. 2019; Hung et al. 2016; C. Li et al. 2015; Yu et al. 2020), however the effect of the channel on migration has varied in the literature, with channel activity supporting migration in some cell types and inhibiting migration in others. Perhaps, a determining factor of the channel’s impact on cell migration is how spatiotemporal localization of the channel is regulated in a given cell type.

Ca^2+^ signals control many aspects of cell migration, including lamellipodial dynamics, traction force generation, rear retraction, focal adhesion turnover and migration directionality (Wei et al. 2012; Tsai et al. 2015; Canales et al. 2019). While mechanically-activated ion channels were proposed to contribute to Ca^2+^ signaling in single cell migration *in vitro* as early as twenty years ago (J. Lee et al. 1999; Doyle, Marganski, and Lee 2004; Wei et al. 2009; Tsai and Meyer 2012; Patkunarajah et al. 2020), many important questions have remained unanswered, including those related to channel identity, functional effects in collective cell migration and physiological contribution during wound healing. We provide evidence for PIEZO1’s involvement in cellular retraction events in single cells as well as in collective cell migration during keratinocyte re-epithelialization. It is well established that retraction during cell migration occurs due to force generated by Myosin II (Cramer 2013; Ridley 2011; Aguilar-Cuenca, Juanes-García, and Vicente-Manzanares 2014), and we previously showed that Myosin II-mediated cellular traction forces elicit localized Piezo1 Ca^2+^ flickers (Ellefsen et al. 2019). Since myosin II activation is enhanced by intracellular Ca^2+^ (Somlyo and Somlyo 2003) these findings suggest that Piezo1 may induce cellular retraction through a feedforward loop between Piezo1 and Myosin II: traction force generation by Myosin II causes Piezo1-mediated Ca^2+^ influx, which in turn may increase Myosin II phosphorylation and force generation through the Ca^2+^-regulated Myosin Light Chain Kinase. Enrichment of Piezo1 in subcellular regions would amplify this effect and result in a localized retraction. Supporting this model, Piezo1-mediated Ca^2+^ events were recently found to elicit retraction of developing endothelial tip cells during vascular pathfinding (Liu et al. 2020) Cell migration involves a complex orchestration of events, including sub-cellular dynamics in which cytoskeletal processes in different compartments of the cell need to be implemented in a precise spatiotemporal order. How this is achieved remains an open question. Our findings suggest that spatiotemporal enrichment dynamics of Piezo1 may play a role in this coordination. More broadly, our findings provide a mechanism by which nanoscale spatial dynamics of Piezo1 channels can control tissue-scale events, a finding with implications beyond wound healing to processes as diverse as development, homeostasis, disease and repair.

## Acknowledgements

We thank Jamison Nourse and Gabriella Bertaccini for technical support, Vivian Leung for help with illustrations, and members of the lab for comments on the manuscript. This work was supported by NIH grants DP2AT010376 and R01NS109810 to M.M.P; NIH grant R01HL143297 to A.P., a James H. Gilliam Fellowship for Advanced Study (GT11549) from the Howard Hughes Medical Institute to M.M.P. and J.R.H., a postdoctoral fellowship from the George Hewitt Foundation for Medical Research to W.-Z.Z., and a seed grant to J.R.H and W.-Z.Z. from the UCI NSF-Simons Center for Multiscale Cell Fate Research (funded by NSF grant DMS1763272 and a Simons Foundation grant 594598). A.P. is an investigator of the Howard Hughes Medical Institute.

## Author Contributions

*Conceptualization,* A.P. S-H.W, M.M.P; *Experimental design,* J.R.H., W-Z.Z., S-H.W, S.M., E.L.E, H.A., M.M.P., A.P.; *Experimentation,* J.R.H., W-Z.Z., S-H.W, E. L. E., S. M., H. A., M. L., M.M.P.; *Live-cell image analysis and interpretation,* J.R.H., M.M.P.; *Formal data analysis,* J.R.H., E. L. E., M.M.P., W-Z.Z., S-H.W, H.A., S.M.; *Visualization,* J.R.H., E.L.E, M.M.P; *Writing—original draft,* J.R.H, M.M.P, E.L.E.; *Writing—review and editing,* M.M.P, E.L.E., J.R.H, A.P., S-H.W., H.A.; all authors read and approved the final draft. *Supervision,* M.M.P and A.P.; *Funding acquisition,* M.M.P and A.P.

## Methods

### Animals

All studies were approved by the Institutional Animal Care and Use Committee of University of California at Irvine and The Scripps Research Institute, as appropriate, and performed in accordance with their guidelines. *Piezo1* LacZ reporter mice (JAX stock 026948) and *Piezo1*-tdTomato reporter mice, expressing a C-terminal fusion of *Piezo1* with tdTomato (*Piezo1*-tdTomato; JAX stock 029214), were generated in a previous study (Ranade et al. 2014). Skin specific *Piezo1-*cKO mice were generated by breeding *Piezo1*^*fl/fl*^ *mice (Cahalan et al. 2015)* (Jax stock 029213) with K14-Cre (The Jackson Laboratory, stock 004782). Skin specific *Piezo1-*GoF mice were generated by breeding *Piezo1*^cx/cx^ mice (Ma et al. 2018) with K14-Cre.

### Keratinocyte isolation

P0-P1 mice were anesthetized with ice prior to decapitation. Bodies were placed in 10% povidone for 1 minute, rinsed with sterile PBS, prior to soaking in 70% ethanol for a further minute, and rinsed again with sterile PBS. Subsequently, the entire dorsal skin was separated from the body. Dorsal skin was left to dissociate in either 0.25% Trypsin/EDTA (Gibco) for 1 hour or 1x dispase solution (Cellntec CnT-DNP-10) at 4°C overnight for 15-18 hours. After incubation, the epidermis was gently separated from the dermis, laid flat, dorsal side down in Accutase (CellnTec CnT-Accutase-100) and incubated for 30 minutes at room temperature. The epidermis was then transferred to a dish of either CnT-02 or CnT-Pr media (CellnTec), supplemented with 10% FBS and 1% Penicillin/Streptomycin. The epidermis was cut into small pieces with scissors prior to agitation on a stir plate for 30 minutes. Cells were then filtered through a 70 μm cell strainer (Falcon) and spun down at 1200 rpm for 5 minutes. The pellet was resuspended in CnT-Pr media (CellnTec) supplemented with ISO-50 (1:1000) (CellnTec) and Gentamicin (50 μg/mL) (ThermoFisher), prior to counting and plating.

### Keratinocyte Culture

For live cell imaging, primary keratinocytes were plated on #1.5 glass-bottom dishes (Mat-Tek Corporation) coated with 10 μg/ml fibronectin (Fisher Scientific, CB-40008A). For monolayer experiments cells were plated at 1.5×10^5^ cells/dish in the 14-mm glass region of dishes. For sparse cell migration experiments and for Ca^2+^ imaging experiments cells were plated at 1.5×10^4^ cells/dish in the 14-mm glass region of dishes. Keratinocytes were imaged following at least 2 days in Cnt-Pr-D (CellnTec) differentiation media.

### Real-time quantitative PCR

After initial keratinocyte isolation and filtering through a 70 μm cell strainer, cells were filtered again through a 40 μm strainer. The filtered solution was spun down and a cell pellet was obtained for RNA isolation. Total RNA was isolated using the RNeasy kit (Qiagen), following which cDNA was synthesized using Superscript III (Invitrogen) and was used for subsequent qPCR experiments (ABI 7900HT fast real time system). qPCR probes (Thermofisher) used were *Piezo1*: Assay ID Mm01241570_g1; ACTGAGAGGATGTTCAGCCAGAATG and Gapdh: Assay ID Mm99999915_g1.

### X-Gal/LacZ staining

Dorsal skin was harvested as described above, cryopreserved in OCT and sectioned into 8 μm thick slices. Skin cryosections were allowed to completely dry prior to being fixed in “fix buffer” composed of 1X PBS, 5 mM EGTA (Sigma cat. # E4378), 2 mM MgCl_2_, 0.2% glutaraldehyde (Sigma cat. # G-7776), pH 7.4 for 15 minutes at room temperature. Next they were washed with “wash buffer” composed of 1X PBS, 2 mM MgCl_2_ twice for 5 minutes each. X-gal staining buffer composed of 1X PBS, 2 mM MgCl_2_, 5 mM potassium ferrocyanide [K_4_Fe(CN)_6_·3H_2_0] (Sigma cat. # P-9287), 5 mM potassium ferricyanide [K_3_Fe(CN)_6_] (Sigma cat. #P-8131), and 1 mg/ ml X-gal [5-bromo-4-chloro-3-indolyl-β-D-galactoside] was made fresh. Then, tissue slides were incubated overnight at 37°C in the “X-gal staining buffer” inside a humidified chamber. The following day, slides were rinsed with 1X PBS and counterstained with Nuclear Fast Red. Slides were then fixed with 4% PFA for longer preservation.

### Immunofluorescence staining

Immunostaining of fixed keratinocytes in Figure 3A was performed as previously described (Ellefsen et al. 2019; Pathak et al. 2014) using primary antibody Goat anti-RFP (Rockland, Cat#200-101-379), 1:200 (5 μg/ml). Secondary antibody used was Donkey anti-goat Alexa Fluor 555 (Invitrogen, Cat#A21432), 1:500. Nuclei were stained by Hoechst (ThermoFisher, Cat#H1399) at 6 μg/ml in PBS. Samples were labeled in Mat-Tek dishes.

For immunostaining of skin sections in Supplementary Figure 3, dorsal skin was prepared and sectioned as for X-Gal staining. Skin cryosections were fixed for 10 minutes in cold acetone, washed twice in 1X PBS prior to blocking for 30 minutes in 10% normal goat serum at room temperature. Primary antibodies used were Rabbit anti-Keratin 14 (Covance, Cat# PRB155P), 1:1000 (1 μg/ml) and Rabbit anti-Keratin 10 (Covance, Cat# PRB-159P), 1:1000 (1 μg/ ml). Secondary antibody used was Goat anti-Rabbit Alexa Fluor 488 (Invitrogen, Cat#A11008), 1:1000. Nuclei were stained by DAPI (Invitrogen, Cat#D1306), 1:50,000. All antibody incubations were performed at room temperature, for 1 hour in 1% BSA in PBS. Slides were mounted in gelvatol containing DAPI.

### Microscopy and image analysis

#### Microscopy

Unless otherwise stated, *in vitro* images were taken using an Olympus IX83-ZDC microscope, equipped with an automated 4-line cellTIRF illuminator. A full enclosure environmental chamber (Tokai Hit) allowed cells to be imaged at 37°C with 5% CO_2_ ensuring optimal cell health during time-lapse experiments. Stage movement was controlled by a programmable motorized stage (ASI) while an Olympus ZDC autofocus control unit allowed for samples to remain in focus throughout imaging periods. The open-source microscopy software μManager was used to control the microscope and acquire images for all except Figure 1A, 4E and Supp Fig 2, 3. Images for Figure 1C, 1E, 2B, 2E, 3, and Supplementary Videos 1, 2, 4, 6, 7, 8, 9, 10 were taken using a PLAPO 60x oil immersion objective with a numerical aperture of 1.45. Images for Figure 3A were taken using a PLAPO 60x oil immersion objective with a numerical aperture of 1.50. Images for Figure 2C, 2D and Supplementary Video 5 were taken using a UPlanSApo 40x dry objective with a numerical aperture of 0.95. Images for Figure 4A, 4B and Supplementary Videos 3, 11 were taken using a UPlanSApo 10x dry objective with a numerical aperture of 0.40. Images for Figure 1C, 1E, 2B, 2C, 2D, 2E, 3A, 3B, 3C, 3D, 3E, 3F, 4A, 4B, and all Supplementary Videos were acquired using a Hamamatsu Flash 4.0 v2+ scientific CMOS camera. Images for Figure 2A were acquired using a Hamamatsu Fusion camera. Images for Supplementary Figure 1B were taken using a Hamamatsu C4742-95-12ER Digital CCD camera.

#### Imaging Piezo1 Ca^2+^ flickers (Fig.1 C-F; Supplementary Videos 1-2)

As described previously, (Ellefsen et al. 2019) TIRF microscopy was used for the detection of Ca^2+^ flickers. Keratinocytes were loaded through the incubation of 2 μM Cal-520 AM (AAT Bioquest Inc.) with 0.04% Pluronic F-127 (ThermoFisher) in phenol red-free DMEM-F12 (Cat#11039047, Gibco) for 30-35 minutes at 37 °C, washed three times, and incubated at room temperature for 10–15 minutes prior to imaging. Cells were imaged at room temperature in a bath solution comprising 148 mM NaCl, 3 mM KCl, 3 mM CaCl_2_, 2 mM MgCl_2_, 8 mM glucose, and 10 mM HEPES (pH adjusted to 7.3 with NaOH, Osmolarity adjusted to 313 mOsm/kg with sucrose). Cal-520 fluorescence was elicited by excitation with a 488 nm laser line and images were acquired at a frame rate of 9.54 frames/second.

#### Piezo1 Ca^2+^ flicker analysis (Fig.1 C-F; Supplementary Videos 1-2)

Ca^2+^ flickers were automatically detected as previously described (Ellefsen et al. 2019) using the detect_puffs plugin (https://github.com/kyleellefsen/detect_puffs) for the open-source image processing program, Flika (https://flika-org.github.io/#). This plugin was used to identify and localize flicker events in recorded videos. Each video is a microscope field of view which contains one or more keratinocytes. To normalize any potential variability in cell number or size between samples, flicker frequency by cell area was computed for each field of view. Cell area was measured by using the ImageJ plugin SIOX: Simple Interactive Object Extraction to create binary masks and compute cell area.

#### Single Cell Tracking Assay (Fig.1 G-J; Supp. Fig. 3)

Primary keratinocytes sparsely seeded on fibronectin-coated glass-bottom dishes were allowed to migrate freely for up to 16.67 hours at 37°C with 5% CO_2_ in bath solution composed of Cnt-Pr-D (CellnTec) culture media with extracellular Ca^2+^ concentration adjusted to 1.2 mM. Timelapse DIC images at multiple microscope fields of view were acquired in each dish of cells at 5 minute intervals for the imaging period. When collecting trajectories we only considered cells which (1) stay within the field of view during the imaging period and (2) did not come into contact with other keratinocytes. The center of the cell body was the tracked position of the cell. The initial positions of cells were manually identified, after which the positions of migrating cells were automatically tracked using the Cell Tracker software (http://www.celltracker.website/index.html) (Piccinini, Kiss, and Horvath 2016). Cell trajectories were logged and exported into Microsoft Excel. Further analysis was subsequently performed using the published open-source algorithm, DiPer (Gorelik and Gautreau 2014) to obtain average instantaneous speed, mean squared displacement (MSD), directionality analysis and trajectory flower plots.

#### Migration dynamics assay (Fig. 2C-E; Supplementary Videos 5-7)

Cells were imaged via DIC microscopy at 37°C with 5% CO_2_, with snapshots taken at 5 second intervals for at least 30 minutes in Cnt-Pr-D culture media (CellnTec) with added 1.2 mM Ca^2+^ and 0.0004% DMSO. After acquiring baseline images, the media was changed and 4 μM Yoda1 was added to the bath solution. After allowing the drug to act for 5-7 minutes, imaging was resumed for at least 55 minutes. Experiments were performed multiple times on independent experiment days. Kymographs were created from representative cells using the ImageJ plugin KymoResliceWide (https://imagej.net/KymoResliceWide). Kymographs were built by taking 1 pixel width lines at regions of interest (ROI) along the cell’s leading edge (Fig. 2).

#### PIEZO1-tdTomato time-lapse imaging (Fig. 2B,3, Supplementary Videos 4, 8-10)

Primary PIEZO1-tdTomato keratinocytes were cultured either sparsely or as confluent monolayers. Monolayers were scratched using a 10 μL pipette tip immediately prior to imaging. Experiments were performed in Cnt-Pr-D (CellnTec) culture media with added 1.2 mM Ca^2+^. Either single cells or, for monolayer scratch experiments, regions along the initial wound edge were marked using a programmable stage and imaged throughout the imaging period as cells migrated to close the wound. PIEZO1-tdTomato channels were illuminated using a 561 nm laser and imaged using TIRF microscopy. TIRF and DIC snapshots at regions of interest were sequentially acquired every 10 (Figures 3 C, D, E, F Supplementary Video 9, 10) or 30 (Figures 3 A, B, Supplementary Figure 5, Supplementary Video 8) minutes over the course of up to 16.67 hours. Time-lapse images were processed from the original recording by subtracting every frame of a movie by (1) the median value z-projection image and then (2) the minimum value z-projection image. KymoResliceWide was then used to construct kymographs by taking the average intensity of the transverse of a 100 pixel wide line drawn at ROIs along the wound edge. ROIs were chosen so that they would capture the wound edge for the entire duration it was present within the microscope field of view. At least one kymograph was generated from each unique microscope field of view. Impartial identification and tracking of periods of channel enrichment was performed using the Kymobutler tool in Wolfram Mathematica (https://gitlab.com/deep-mirror/kymobutler). Only puncta located at the wound edge that were successfully tracked for at least 70 minutes were collected for further analyses. The pixel classification function of the computer vision software, Ilastik (https://www.ilastik.org/) (Berg et al. 2019) was used to classify DIC images and create binarized movies used for generating binarized kymographs in Figure 3 E,F.

#### Wound edge dynamics assay (Fig. 4A, 4B, Supplementary Video 11)

Primary keratinocytes were densely seeded and cultured to form monolayers. Scratch wounds were generated by scratching monolayers with a 10 μl pipette tip immediately prior to imaging. Dishes were washed 3x with cell culture media to remove cell debris. Yoda1 (4 μM) or equivalent concentration of DMSO control was added to the bath media immediately prior to imaging. DIC snapshots were taken every 5 minutes at ROIs along the wound edge for at least 9 hours. Representative kymographs were created using 1 pixel wide ROIs using KymoResliceWide. Binarized movies generated by using the ImageJ plugin Trainable Weka Segmentation (https://imagej.net/Trainable_Weka_Segmentation) were used to create binarized kymographs.

#### In vitro wound healing assay (Fig. 4C-D)

Primary keratinocytes were seeded in a two-well silicone insert in a 35 mm dish (ibidi, 81176). Cells were cultured until a monolayer confluence was reached. Subsequently, the insert was removed from the dish to create a reproducible 500 μm cell-free gap. The dishes were washed with cell culture medium to remove floating cells and cell debris. For pharmacology experiments, Yoda1 (4 μM) or equivalent concentration of DMSO control was added into the dishes immediately prior to imaging. Dishes were imaged by phase-contrast microscopy at 37°C (5% CO_2_) for indicated time points.

### In vivo wound healing assay

Adult (3-4 month) male and female mice were anaesthetized with isoflurane and placed on a heated blanket. The dorsal hair was shaved and further removed by hair-removal cream. Two full-thickness wounds were created in the upper dorsal skin using a 4 mm wide dermal biopsy punch (Integra LifeSciences Corporation). Wounded areas were patched with medical dressing, Tegaderm (3M). Wound sizes were measured with a scale loupe (Peak Optics, #1975) at Day 6 to compare healing progress. Both the short (dS) and long (dL) diameters of the oval-shaped wounds were measured and used to calculate an overall wound area using the equation: dS × dL × ϖ.

### Statistical Analysis

All of the data are presented as the mean ± SEM. Sample sizes are indicated in corresponding figure legends. OriginPro 2020 (OriginLab Corporation) was used for statistical analysis and generating plots. *P* values and statistical tests used are indicated in figure legends. Unless otherwise stated, a two-sample *t*-test was used where data were modeled by a normal distribution and the nonparametric Kolmogorov–Smirnov test was used in the case of non-normal distributions.

### Code Availability

No custom code was generated for performing data analysis.

