## Supplementary Material for "Spatiotemporal dynamics of PIEZO1 localization controls keratinocyte migration during wound healing"

### Supplementary Figures

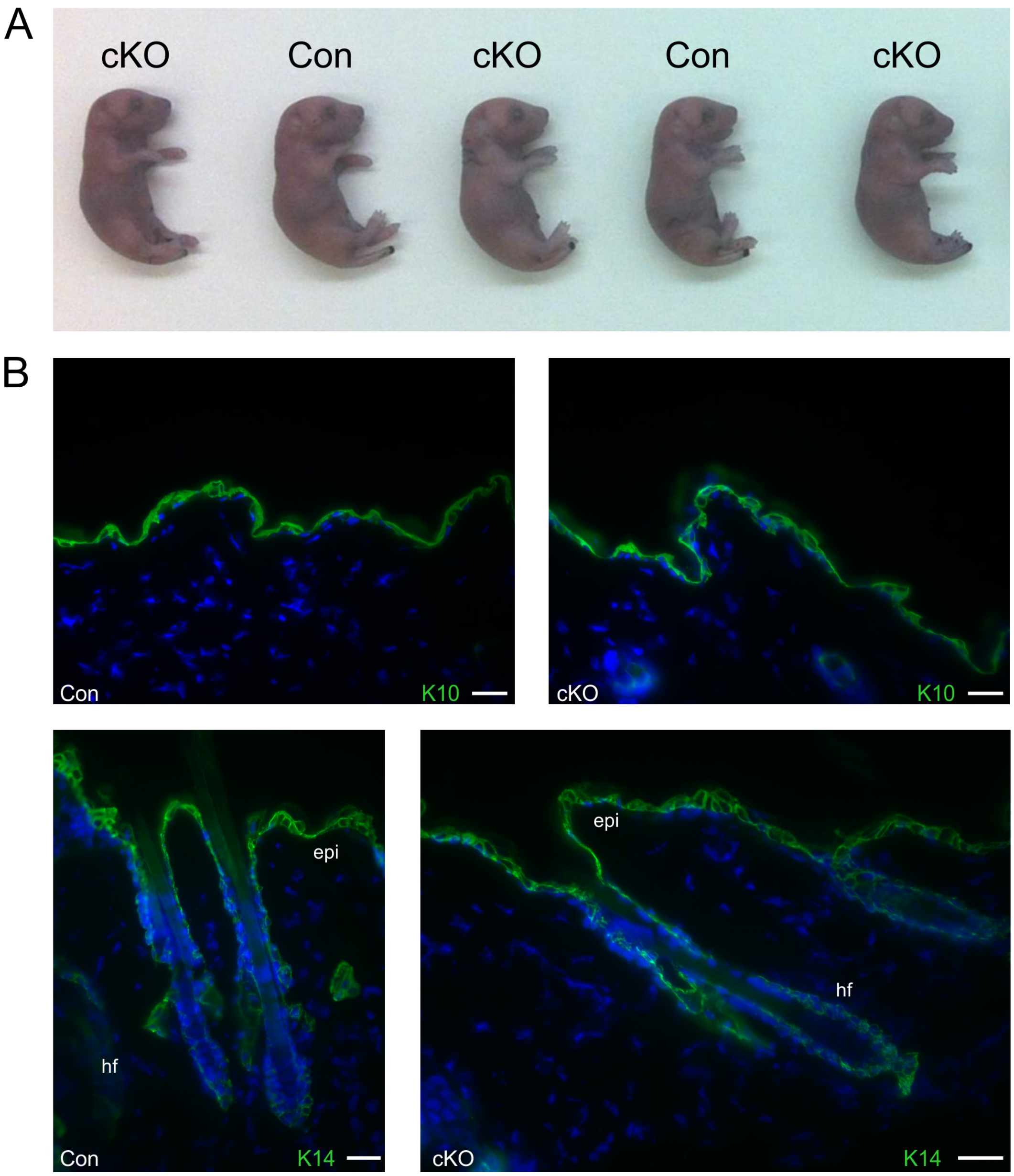


**Supplementary Figure 1. *Piezo1*-cKO mice develop normally. (A)** Images of P2 pups from *Krt14Cre;Piezo1*$^{fl/fl}$ (cKO) and Cre- (Con) littermates illustrating normal development of neonatal pups. **(B)** Staining of keratin markers (Green), K10 (*top)* and K14 (*bottom)*, on skin sections derived from adult *Krt14Cre;Piezo1*$^{fl/fl}$ (cKO) and Cre- (Con) littermates, highlighting normal patterns of expression in Control (*left*) and cKO (*right*). epi, epidermis. hf, hair follicle. Scale bar = 20 µm.

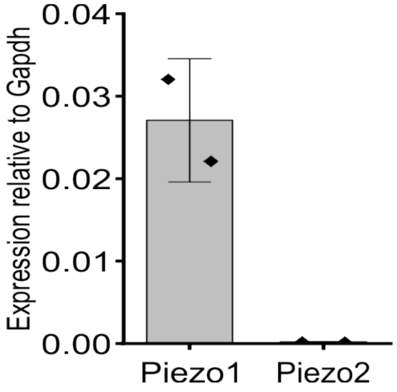


**Supplementary Figure 2. *Piezo1* is the primary Piezo channel in mouse keratinocytes.** qRT-PCR from primary neonatal keratinocytes of *Piezo1* and *Piezo2* mRNA expression relative to *Gapdh*.

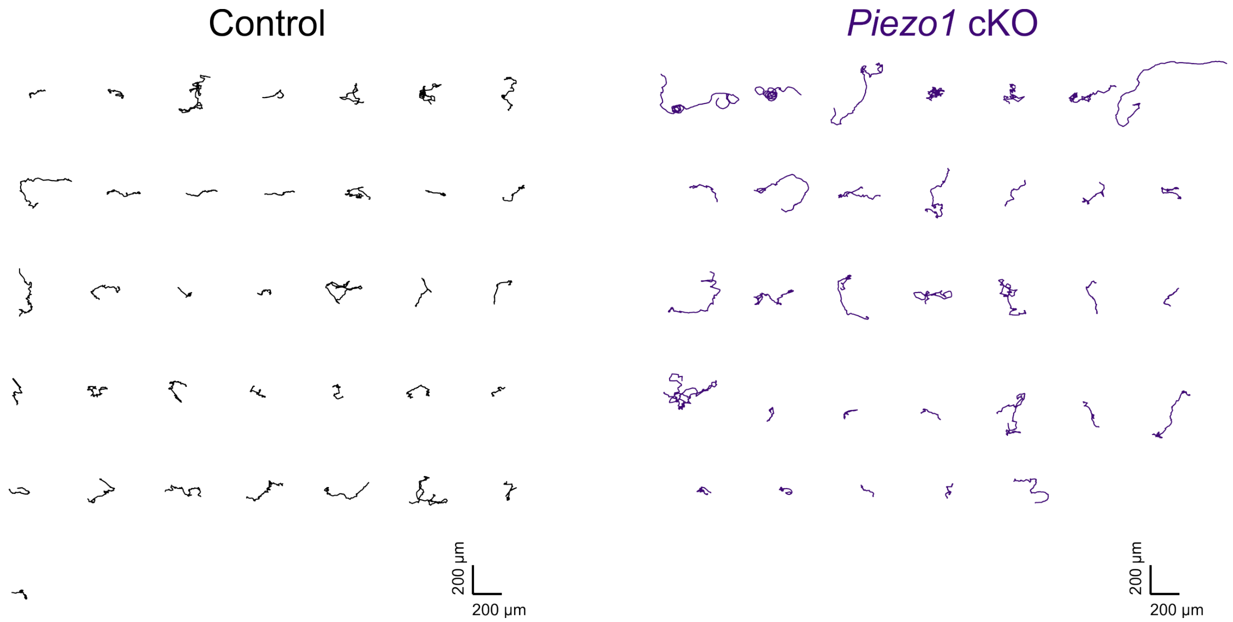


**Supplementary Figure 3. *Piezo1*-cKO keratinocytes migrate further. (A)** Individual trajectories seen in Figure 2 from *Krt14Cre;Piezo1$^{fl/fl}$* (cKO) mice and littermate controls.

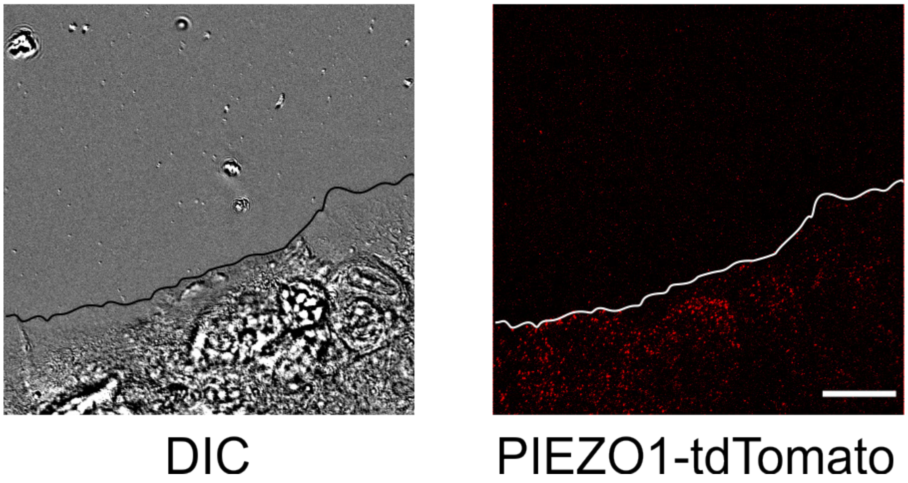


**Supplementary Figure 4. Absence of PIEZO1-tdTomato enrichment at wound edge immediately after scratch wound generation.** Representative DIC (*left*) and TIRF (*right)* images illustrating the location of PIEZO1-tdTomato protein immediately after wounding in a keratinocyte monolayer in an *in vitro* scratch assay. Black (*left*) and white (*right*) lines denote the cell boundary. This is the same field of view as in Fig. 4C. Note the absence of PIEZO1-tdTomato enrichment at the wound edge as compared to 11 hours later in Fig. 4C. Scale Bar = 20 μm.

### Description of Supplementary Videos

**Supplementary Video 1. Piezo1 $Ca^{2+}$ flickers are reduced in *Piezo1* knockout keratinocytes.** The video shows an $F/F_0$ ratio movie of $Ca^{2+}$ flickers in control (*left)* and *Piezo1*-cKO (*right)* keratinocytes. Related to Fig. 1C-E.

**Supplementary Video 2. Piezo1 $Ca^{2+}$ flickers are increased in *Piezo1* Gain-of-Function keratinocytes.** The video shows an $F/F_0$ ratio movie of $Ca^{2+}$ flickers in control (*left*) and *Piezo1*-GoF (*right*) keratinocytes. Related to Fig. 1F-H.

**Supplementary Video 3. *Piezo1* knockout affects keratinocyte motility.** The video shows representative DIC time-lapse images of single migrating control (*left)* and *Piezo1*-cKO (*right)* keratinocytes. Related to Fig. 2A.

**Supplementary Video 4. PIEZO1 agonist Yoda1 increases retraction in keratinocytes.** A representative DIC time-lapse movie showing lamellipodial dynamics in response to treatment with 4 µM Yoda1. Imaging was performed in control medium for 30 min, after which Yoda1 was added, the cell was incubated for 5-7 minutes, and imaging was resumed. Related to Fig. 3A, B.

**Supplementary Video 5. PIEZO1 agonist Yoda1 can cause widespread retraction in keratinocytes.** A representative DIC time-lapse movie showing marked cellular retraction in response to treatment with 4 µM Yoda1. Imaging was performed in control medium for 55 minutes, after which Yoda1 was added, the cell was incubated for 5-7 minutes, and imaging was resumed. Related to Fig. 3.

**Supplementary Video 6. Yoda1 does not increase retraction in *Piezo1*-knockout keratinocytes.** A representative DIC time-lapse video showing lamellipodia dynamics of *Piezo1*-cKO keratinocytes in control medium for 55 min, after which 4 µM Yoda1 was added to imaging solution, the cell was incubated for 5-7 minutes, and imaging was resumed. Related to Fig. 3C.

**Supplementary Video 7. PIEZO1 agonist Yoda1 inhibits migration of keratinocyte monolayers.** A representative DIC time-lapse series showing healing monolayers of keratinocytes during an overnight *in vivo* scratch assay under control (*left)* and +4 µM Yoda1 (*right)* conditions. Related to Fig. 3D, E.

**Supplementary Video 8. PIEZO1-tdTomato is enriched at the rear of individually migrating keratinocytes.** Representative video shows DIC, TIRF (for PIEZO1-tdTomato) and DIC + TIRF merged videos from an individually migrating PIEZO1-tdTomato keratinocyte. Related to Fig. 4B.

**Supplementary Video 9. Lack of PIEZO1-tdTomato enrichment in advancing monolayers.** Representative video showing a time-lapse of DIC, TIRF (PIEZO1-tdTomato) and DIC + TIRF merged images from a monolayer of PIEZO1-tdTomato keratinocytes corresponding to Fig. 4E. Note lack of enrichment at the monolayer edge throughout video, as the wound edge advances. Related to Fig. 4E, G.

**Supplementary Video 10. PIEZO1-tdTomato enrichment becomes enriched at the front of keratinocyte monolayers and elicits retraction.** Representative video showing time-lapse of the DIC, TIRF (PIEZO1-tdTomato) and DIC + TIRF merged images from a monolayer of

PIEZO1-tdTomato keratinocytes corresponding to Fig. 4D. Arrows denote period of channel enrichment starting at ~7.5 hours. Note the PIEZO1-tdTomato enrichment during periods of retraction and absence of channel enrichment during periods of protrusion. Related to Fig. 4D, F.

**Supplementary Video 11. Persistent PIEZO1-tdTomato enrichment at the wound edge elicits sustained retraction.** Representative video shows time-lapse of DIC, TIRF (PIEZO1-tdTomato) and DIC + TIRF merged images of a monolayer of PIEZO1-tdTomato keratinocytes. Note the enrichment of PIEZO1-tdTomato at the wound edge and the ensuing retraction. Related to Fig. 4H.
